# Recurrent Neoantigens in Colorectal Cancer as Potential Immunotherapy Targets

**DOI:** 10.1101/840918

**Authors:** Chao Chen, Songming Liu, Bo Li

## Abstract

This study was aimed to investigate the mutations in colorectal cancer (CRC) for recurrent neoantigen identification. A total of 1,779 samples with whole exome sequencing (WES) data were obtained from 7 published CRC cohorts. Common HLA genotypes were used to predict the probability of neoantigens at high frequency mutants in the dataset. Based on the WES data, we not only obtained the most comprehensive CRC mutation landscape so far, but also found 1550 mutation sites which could be identified in at least 5 or more patients, including *KRAS* G12D (8%), *KRAS* G12V (5.8%), *PIK3CA* E545K (3.5%), *PIK3CA* H1047R (2.5%) and *BMPR2* N583Tfs*44 (2.8%). These mutations can also be recognized by multiple common HLA molecules as potential ‘public’ neoantigens. Many of these mutations also have high mutation rates in metastatic pan-cancers, suggesting their value as therapeutic targets in different cancer types. Overall, our analysis provides recurrent neoantigens as potential cancer immunotherapy targets.

## 1. Introduction

Colorectal cancer (CRC) is the third most common malignancy in the world and the second leading cause of cancer-related mortality[1, 2]. Traditional treatments, such as surgery, chemotherapy and radiation, have been important in prolonging patients’ survival, but for patients with advanced CRC, especially those with metastatic disease, these treatments are limited and often intolerant[3].

In recent years, immunotherapy, including immune checkpoint inhibitors (ICIs), cancer vaccines and neoantigen-based tumor-infiltrating lymphocytes (TILs), has played an increasingly important role in cancer therapy[4]. Certain CRC patients with microsatellite instability (MSI) could potentially benefit from ICIs treatment[5]. However, not all CRC patients with MSI show clinical efficacy in ICIs treatment. Neoantigen-based immunotherapy is complementary to ICIs since it has no specific requirement for patient’s MSI status nor tumor mutation burden (TMB). Current studies on CRC genomics mainly focus on the dissection of tumor occurrence and metastasis mechanisms, and rarely analyze into tumor-specific neoantigens [6–11]. By integrating the mutation data of already existing CRC cohorts and combining the common HLA genotypes in this population[12, 13], our study is expected to find the common neoantigens in CRC patients and facilitate further development of off-the-shelf neoantigen-based immunotherapy.

## 2. Materials and Methods

### 2.1. Genomic Data of CRC

This study was approved by the Institutional Review Board on Bioethics and Biosafety of BGI group. All somatic mutations, including single nucleotide variants (SNVs) and short insertion/deletion (indels), were downloaded from the latest publications (Table 1 and Supplementary Table S1), which represent seven geographically diverse study groups involving 1779 CRC patients. Since all data used in this study were from public databases with informed consent from participants in the original genome study, no additional informed consent was required.

**Table 1:** Summary of clinical information of CRC cohort, including patients from seven studies and 1779 CRC samples.

| Characteristic | Baylor<br>(n=110) | Beijing<br>(n=98) | COCA-CN<br>(n=321) | Genetech<br>(n=74) | Harvard<br>(n=619) | TCGA<br>(n=528) | Texas<br>(n=29) | Total<br>(n=1779) |
| --- | --- | --- | --- | --- | --- | --- | --- | --- |
| **Age (years)** |  |  |  |  |  |  |  |  |
| <60 | 38 (34.5%) | 48 (49.0%) | 155 (48.3%) | 0 (0%) | 67 (10.8%) | 158 (29.9%) | 0 (0%) | 466 (26.2%) |
| >=60 | 70 (63.6%) | 50 (51.0%) | 166 (51.7%) | 0 (0%) | 550 (88.9%) | 366 (69.3%) | 0 (0%) | 1202 (67.6%) |
| Unknown | 2 (1.8%) | 0 (0%) | 0 (0%) | 74 (100%) | 2 (0.3%) | 4 (0.8%) | 29 (100%) | 111 (6.2%) |
| **Sex** |  |  |  |  |  |  |  |  |
| Female | 65 (59.1%) | 50 (51.0%) | 127 (39.6%) | 0 (0%) | 380 (61.4%) | 253 (47.9%) | 15 (51.7%) | 890 (50.0%) |
| Male | 45 (40.9%) | 48 (49.0%) | 194 (60.4%) | 0 (0%) | 239 (38.6%) | 273 (51.7%) | 14 (48.3%) | 813 (45.7%) |
| Unknown | 0 (0%) | 0 (0%) | 0 (0%) | 74 (100%) | 0 (0%) | 2 (0.4%) | 0 (0%) | 76 (4.3%) |
| **Stage** |  |  |  |  |  |  |  |  |
| I | 12 (10.9%) | 10 (10.2%) | 40 (12.5%) | 0 (0%) | 152 (24.6%) | 94 (17.8%) | 0 (0%) | 308 (17.3%) |
| II | 42 (38.2%) | 44 (44.9%) | 94 (29.3%) | 0 (0%) | 187 (30.2%) | 196 (37.1%) | 0 (0%) | 563 (31.6%) |
| III | 48 (43.6%) | 39 (39.8%) | 130 (40.5%) | 0 (0%) | 159 (25.7%) | 150 (28.4%) | 1 (3.4%) | 527 (29.6%) |
| IV | 8 (7.3%) | 4 (4.1%) | 56 (17.4%) | 0 (0%) | 65 (10.5%) | 69 (13.1%) | 28 (96.6%) | 230 (12.9%) |
| Unknown | 0 (0%) | 1 (1.0%) | 1 (0.3%) | 74 (100%) | 56 (9.0%) | 19 (3.6%) | 0 (0%) | 151 (8.5%) |
| **MSI.Status** |  |  |  |  |  |  |  |  |
| MSI-H | 24 (21.8%) | 8 (8.2%) | 0 (0%) | 15 (20.3%) | 91 (14.7%) | 65 (12.3%) | 0 (0%) | 203 (11.4%) |
| MSI-L | 0 (0%) | 0 (0%) | 0 (0%) | 0 (0%) | 0 (0%) | 77 (14.6%) | 0 (0%) | 77 (4.3%) |
| MSS | 81 (73.6%) | 32 (32.7%) | 0 (0%) | 59 (79.7%) | 438 (70.8%) | 346 (65.5%) | 29 (100%) | 985 (55.4%) |
| Unknown | 5 (4.5%) | 58 (59.2%) | 321 (100%) | 0 (0%) | 90 (14.5%) | 40 (7.6%) | 0 (0%) | 514 (28.9%) |

### 2.2. Pipeline for Neoantigen Prediction

For neoantigen prediction, a total of 43 HLA genotypes were selected with frequencies greater than 5% in the Chinese or TCGA cohort. Mutations, which include 76 non-silent somatic SNVs and 973 indels, could be identified in at least 5 patients. All these mutations were then used for neoantigen prediction by NetMHC[14], NetMHCpan[15], PickPocket[16], PSSMHCpan[17], and SMM[18]. According to our previous research[19], neoantigen peptides need to meet the following three criteria: (1) Between 8-11 mers length; (2) Affinity IC50 < 500nM on at least two tools; (3) Mutant (MT) peptides affinity of score lower than the wild type (WT).

### 2.3. Statistical Analysis

The statistical analysis was done in R-studio and the mutation analysis and drawing were done with the maftools package[20]. If no special instructions were given, P < 0.05 was considered significant.

## 3. Results and Discussion

### 3.1. The Integrated Mutation Landscape of CRC Patients

The mutation status of all samples is shown in **Figure 1 and Figure S1**. In general, missense mutation is the most main type of mutations. At the base substitution level, C>T is the dominant mutant form, followed by C>A (**Figure S2**), which is consistent with TCGA and previous reports[21, 22]. The median number of mutations in each sample was 110, among which *APC*, *MUC16, TP53, SYNE1, KRAS* and *PIK3CA* were the most frequently mutated genes. Apart from the samples without MSI information, there are 203 MSI-H samples in the combined cohort (**Table 1**), accounting for 11.4%. The mutation load of MSI samples is higher than the MSS samples with more indel mutations (**Figure S3-S4**).

**Figure 1.**
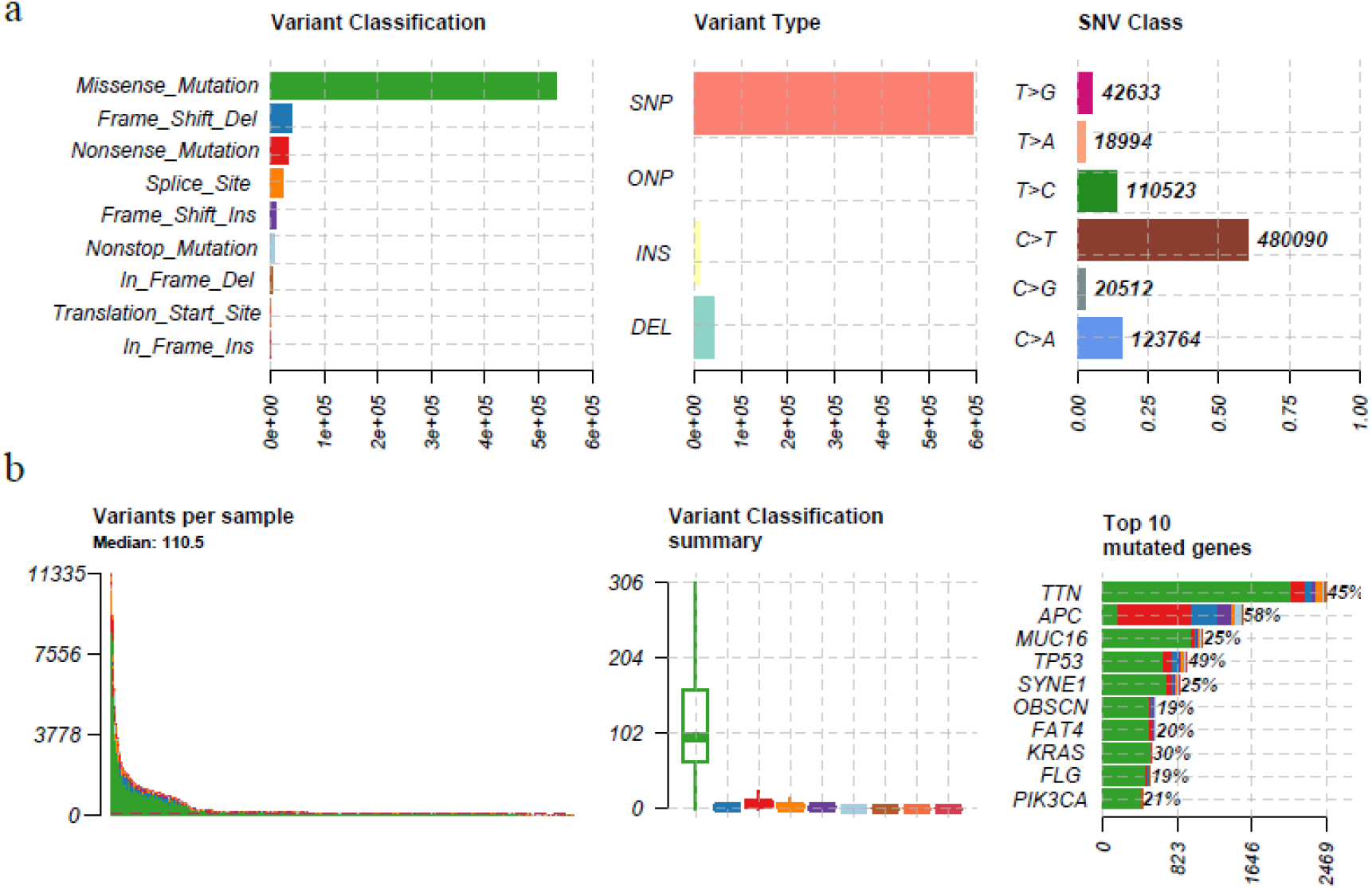
The mutation landscape in CRC cohort. (a) from left to right, counts of each variant classification, counts of each variant type, counts of each SNV class. (b) from left to right, variants number per sample, variant classification and top 10 significantly mutated genes.

Interestingly, through the integration analysis of mutation pattern in MSS samples, we found the mutation of four genes, including *TENM1*, *SOX9*, *PIK3CA* and *KRAS*, and the mutation of TP53 gene, were mutually exclusive (Fisher’s exact test, P<0.05, **Figure S5**). This different mutation pattern may suggest that the carcinogenic mechanisms are different in CRC patients carrying mutations in these four genes and in those carrying *TP53* mutation. Correspondingly, there is no such mutual exclusion effect in MSI samples (**Figure S6**).

There are many hot spot mutations in CRC samples. Of these, 1550 recurrent mutations could be identified in at least 5 patients, with 476 SNVs and 974 indels, respectively. Previous studies have shown that common neoantigens in cancers could be used as potential immunotherapy targets[19, 23]. Therefore, in order to find out whether there are common neoantigens in CRC populations, we used these mutations in downstream analyses to predict tumour-specific neoantigens.

### 3.2. Neoantigens Shared among CRC Patients

Due to the difference in the frequency of HLA in different populations, in order to search for ‘public’ neoantigens in CRC populations, we selected high-frequency HLA in Chinese (HLA frequency >5% in Han Chinese [13]) and high-frequency HLA in Americans (HLA frequency >5% in TCGA [24]) for neoantigen analysis. Finally, a total of 43 HLA alleles were used for neoantigen prediction (**Table S2**).

We detected 274 SNV derived neoantigens and 1269 indel derived neoantigens (**Table S3-4**). Each SNV usually produces 1-2 high-affinity peptides, while each indel can produce multiple high-affinity peptides. The top ten high frequency mutation sites and neoantigen peptides in SNV and indel are shown in **Table 2** and **Table 3**, respectively. In terms of SNV, mutations of *KRAS, PIK3CA, PCBP1* and *CHEK2* can produce 10 neoantigens with the highest frequency. In terms of indel, although the mutation frequency is not as high as SNV, generally one indel can produce about 5-10 neoantigen peptides.

**Table 2.**
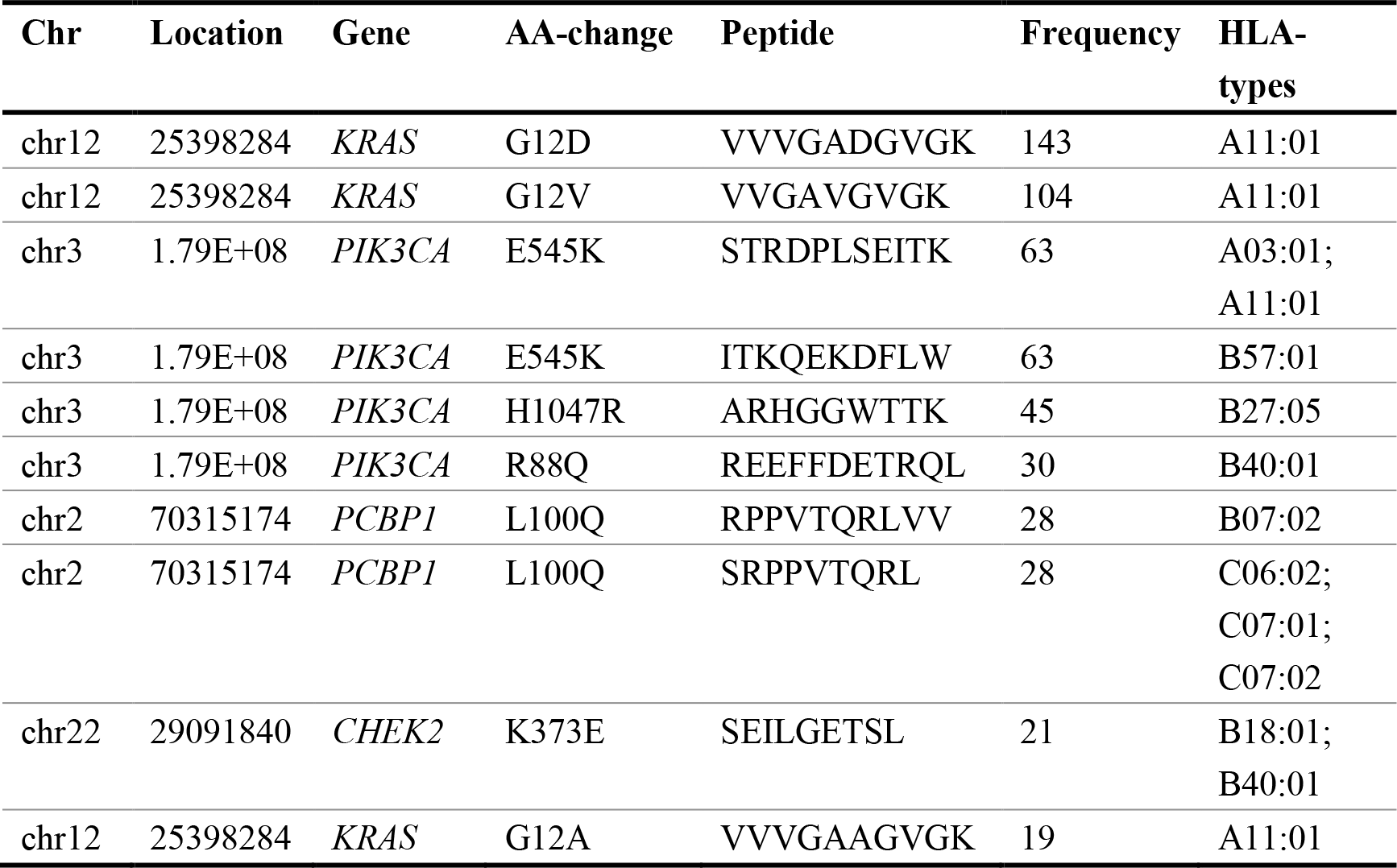
Top ten SNVs and the corresponding neoantigens in CRC cohort.

**Table 3.** Top ten indels and the corresponding frequency in CRC cohort.

| Chr | Location | Gene | AA-change | Frequency |
| --- | --- | --- | --- | --- |
| chr2 | 203420130 | *BMPR2* | N583Tfs*44 | 50 |
| chr10 | 890939 | *LARP4B* | T163Hfs*47 | 37 |
| chr1 | 1290110 | *MXRA8* | R301Gfs*107 | 31 |
| chr18 | 34205516 | *FHOD3* | S336Vfs*138 | 29 |
| chr15 | 45003781 | *B2M* | L15Ffs*41 | 28 |
| chr12 | 110019240 | *MVK* | A141Rfs*18 | 27 |
| chr3 | 168833257 | *MECOM* | G614Efs*30 | 26 |
| chr22 | 20130522 | *ZDHHC8* | T459Rfs*177 | 20 |
| chr8 | 103289349 | *UBR5* | E2121Kfs*28 | 20 |
| chr6 | 158508009 | *SYNJ2* | P1113Lfs*5 | 19 |

### 3.3. Comparison of Neoantigens in Different Subtypes of CRC

By comparing the neoantigen profiles of different subtypes of CRC, we found that there were more SNV and indel derived neoantigens in MSI-H CRC than in MSS patients (Fisher’s exact test, P<0.01, **Figure 2a**). The 1269 indel related neoantigens can cover 86.7% of patients with MSI-H colorectal cancer, but only 3.9% of MSS patients were covered, indicating that indel is the main source neoantigens of MSI-H CRC. SNV derived neoantigens can cover 41% of MSS patients and 66% of MSI patients, respectively.

**Figure 2.**
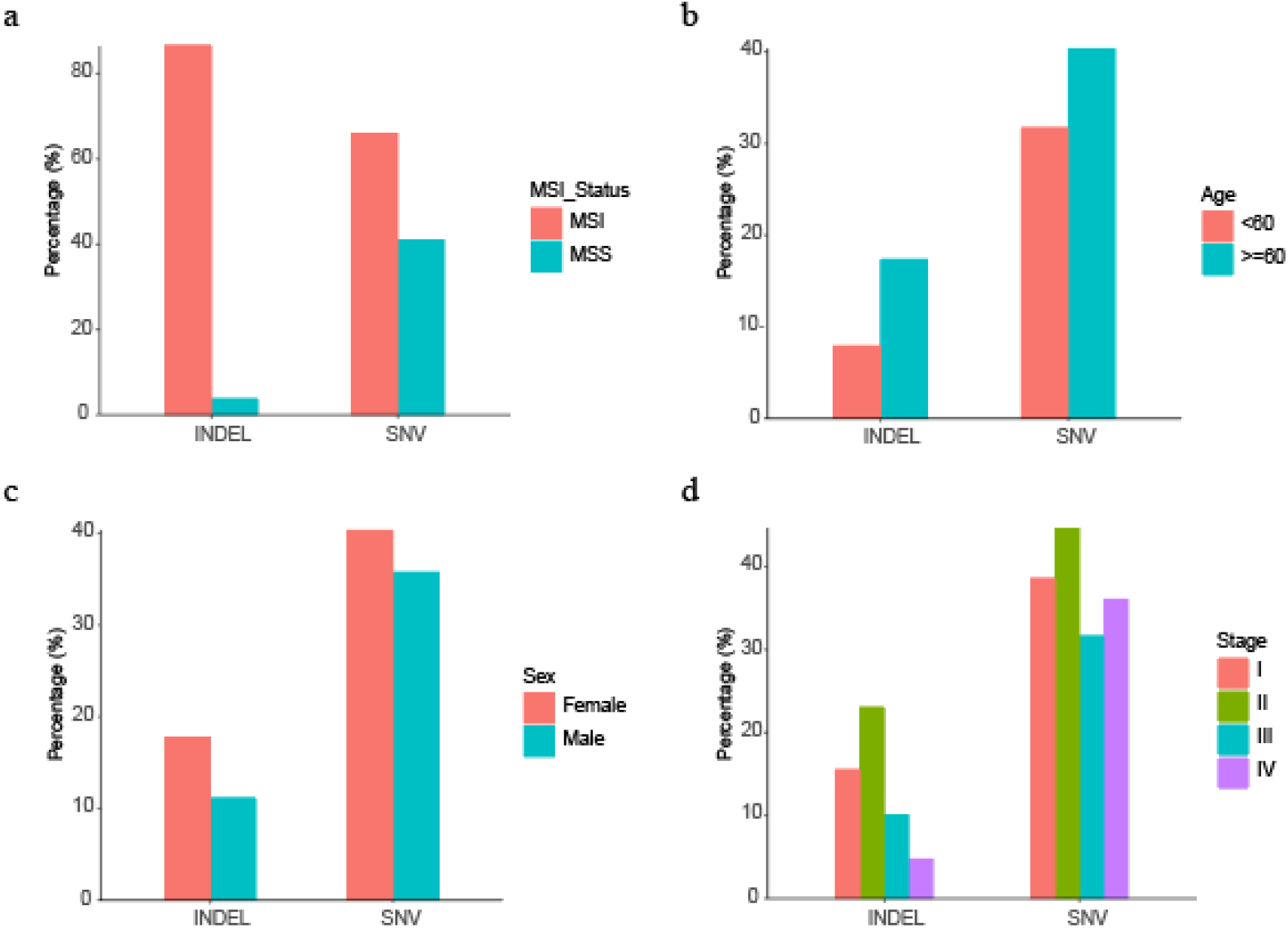
The comparation of neoantigens between different subgroups. (a) between different MSI type; (b) between age≧60 and age <60 groups; (c) between female and male groups; (d) between different stage. These analyses excluded patients with unknown subtypes.

In the comparison of age classification, patients older than 60 carry more neoantigens, with both SNV derived and indel derived neoantigens (Fisher’s exact test, P<0.01, **Figure 2b**). Women tend to carry more neoantigens than men, especially those derived from indels. In terms of cancer stage, Stage II CRC patients carry the most abundant neoantigens, which may be related to the mutation load of the corresponding subgroup (**Figure 2c-d**).

### 3.4. Hotspot Mutation-related Neoantigens that May be A Potential Source of Immunotherapy Target in CRC and Pan-cancer

To further investigate the potential significance of these high frequency neoantigens, we focused on sites with the highest mutation frequency, including G12D (*KRAS*), G12V (*KRAS*), E545K (*PIK3CA*), and H1047R (*PIK3CA*), because these sites not only produce recurrent neoantigens, but also have a higher mutation frequency in CRC cohort (**Figure 3a-b**).

**Figure 3.**
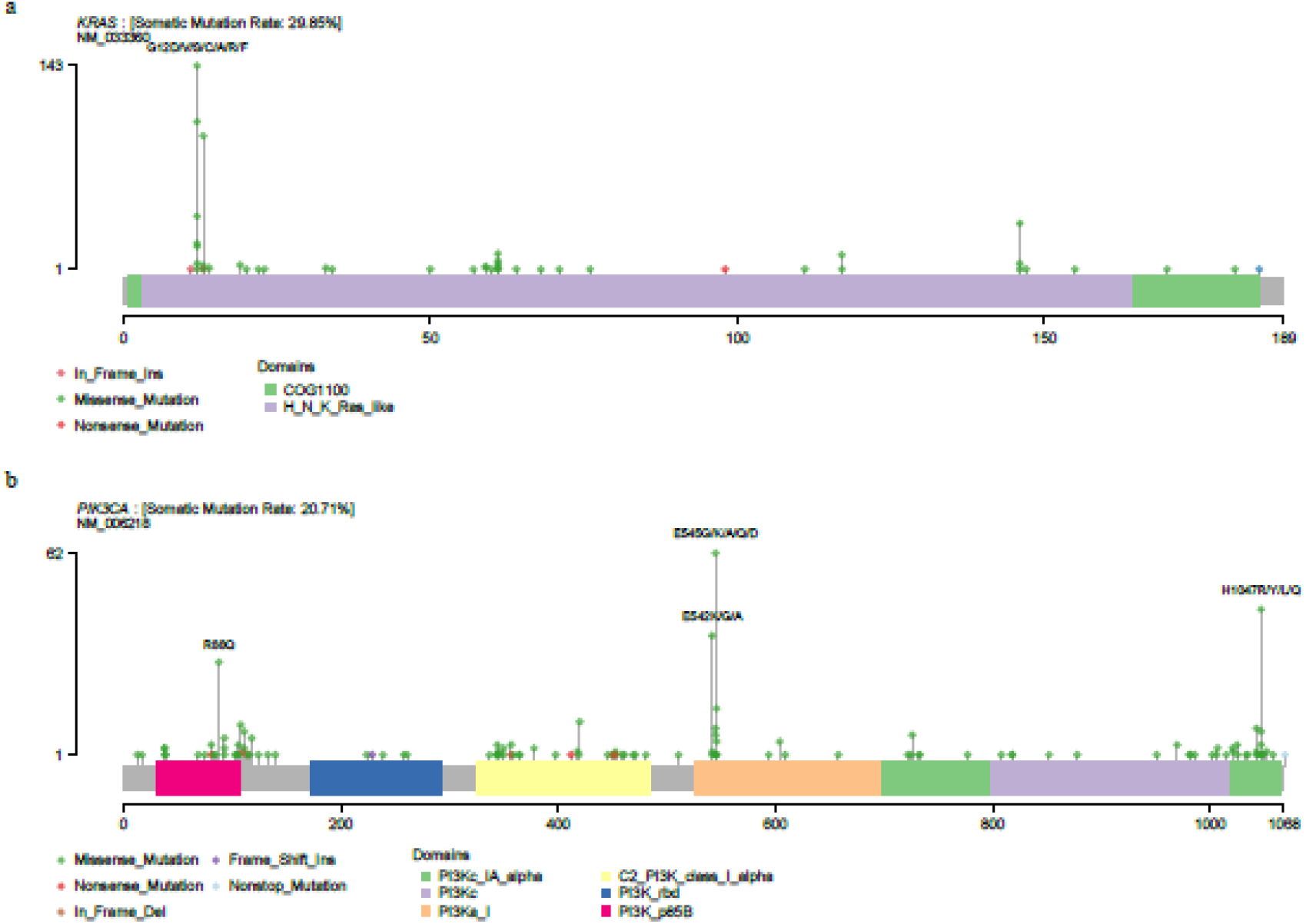
Mutational spectrum of KRAS (a) and PIK3CA (b) in 1179 CRC patients.

*KRAS* Gly12 (including G12V, G12C, and G12D) is a classic cancer mutation that occurs more than 20% in the metastatic pancreatic and appendiceal cancers [24]. Both G12D and G12V are highly mutated in multiple metastatic cancers, including Endometrial, CRC, and non-small cell lung cancer (**Figure 4a-b**). Witkiewicz et al. have reported the high frequency of *KRAS* G12D mutations in pancreatic cancer [23,25]. Liang et al. also demonstrated by mass spectrometry in 2019 that *KRAS* G12V mutated neoantigen can be presented by HLA-A11:01 cell lines[25]. And as far as we know, clinical trials for *KRAS* G12V mutations in patients with HLA-A11:01 cancer are already under way (NCT03190941).

**Figure 4.**
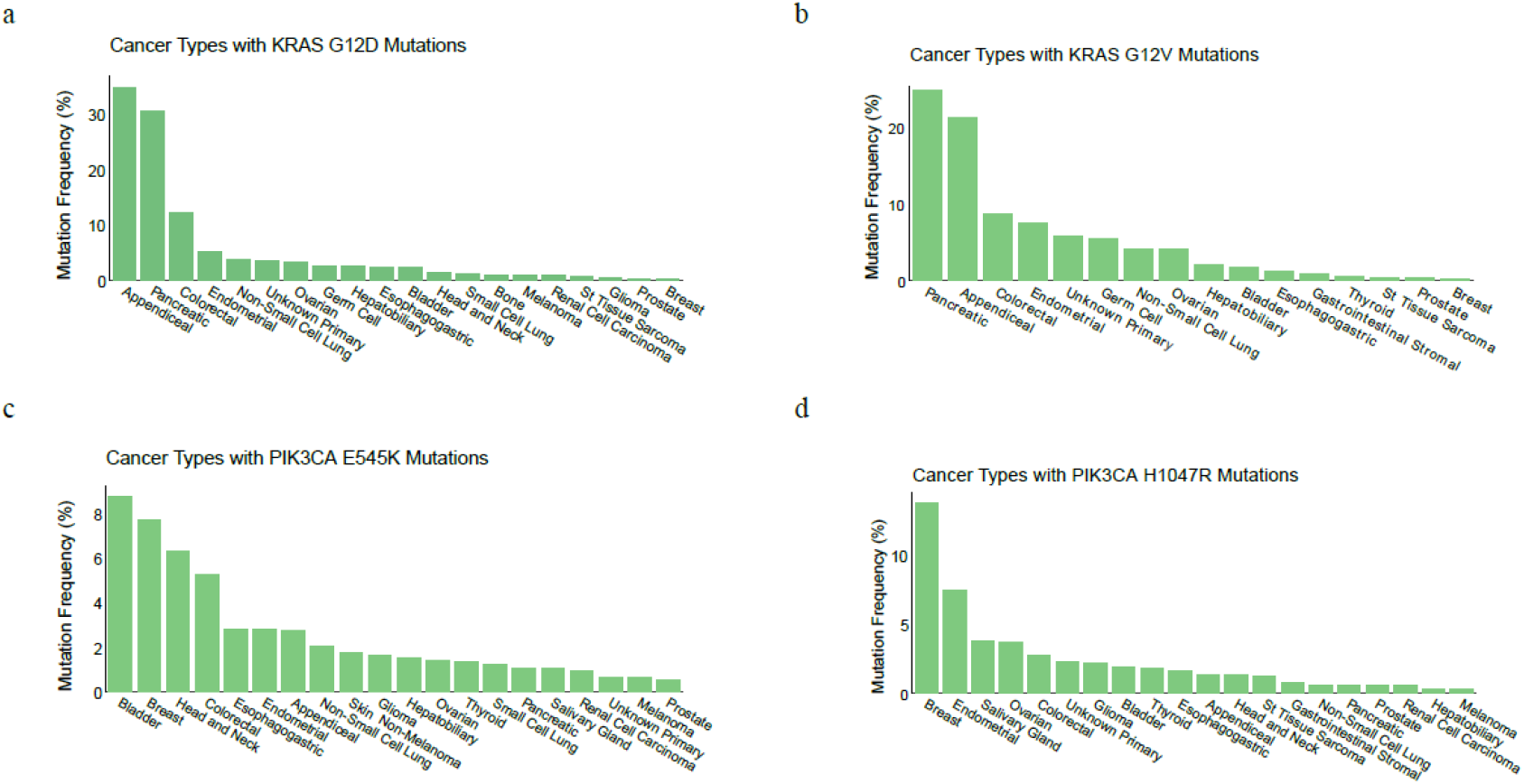
Mutation frequency in MSK-IMPACT cohorts. *KRAS* G12D (a), *KRAS* G12V (b), *PIK3CA* E545K (c), *PIK3CA* H1047R (d) in MSK-IMPACT pan-cancer cohorts.

*PIK3CA* is one of the hot driving genes in gastrointestinal malignancies[26, 27]. *PIK3CA* E545K is a hot spot mutation in breast cancer and has corresponding first-line drugs (Alpelisib and Fulvestrant)[28]. In addition to breast cancer, this mutation could be found in more than 5% bladder, Head and Neck, and colorectal cancer patients (**Figure 4c**). *PIK3CA* H1047R is most commonly found in breast cancer[29] and is also frequently mutated among multiple tumor types in the MSK-IMPACT metastatic cancers (**Figure 4d**). Our previous studies have shown that this mutation can be presented by multiple HLA molecules (e.g., HLA-C07:02, HLA-C 07:01, HLA-A30:01, HLA-B58:01) and can be a potential neoantigen in patients with gastric cancer[19]. Combined with the results of this study, it is suggested that this mutation can be used as an important therapeutic target for patients with gastrointestinal tumors.

## 4. Conclusion

Based on the analysis of the published WES data of CRC, the most complete mutation landscape of CRC was constructed. We selected HLA subtypes with high frequency in Chinese and TCGA cohort to predict the common neoantigens in the population. The high frequency mutations, including *KRAS* G12D (8%), *KRAS* G12V (5.8%), *PIK3CA* E545K (3.5%), *PIK3CA* H1047R and *BMPR2* N583Tfs*44, can be recognized by many HLA genes, such as HLA-A1101, HLA-A03:01 and HLA-B57:01. These HLA genotypes are the main HLA subtypes in Chinese and Americans, indicating the broad spectrum of the neoantigens we identified. In conclusion, we have found a series of public neoantigens for CRC, which provide important resources for immunotherapy of CRC in the future.

## Acknowledgements

We thank for the support of the National Natural Science Foundation of China (No.81702826), the Science, Technology and Innovation Commission of Shenzhen Municipality under grant No. JSGG20170824152728492, the Science, Technology and Innovation Commission of Shenzhen Municipality under grant No. JCYJ20170303151334808.

